# Transcriptional Activation of Arabidopsis Zygotes Is Required for Their Initial Division

**DOI:** 10.1101/679035

**Authors:** Ping Kao, Michael Nodine

## Abstract

Commonly referred to as the maternal-to-zygotic transition, the shift of developmental control from maternal-to-zygotic genomes is a key event during animal and plant embryogenesis. Together with the degradation of parental gene products, the increased transcriptional activities of the zygotic genome remodels the early embryonic transcriptome during this transition. Although evidence from multiple flowering plants suggests that zygotes become transcriptionally active soon after fertilization, the timing and developmental requirements of zygotic genome activation in *Arabidopsis thaliana* (Arabidopsis) remained a matter of debate until recently. In this report, we optimized an expansion microscopy technique for robust immunostaining of Arabidopsis ovules and seeds. This enabled the detection of marks indicative of active transcription in zygotes before the first cell division. Moreover, we employed a live-imaging culture system together with transcriptional inhibitors to demonstrate that such active transcription is required in zygotes. Our results indicate that zygotic genome activation occurs soon after fertilization and is physiologically required prior to the initial zygotic division in Arabidopsis.

## INTRODUCTION

The transition of developmental control from parental-to-zygotic genomes is a pivotal event during animal and plant development. In animals, maternally inherited gene products regulate early embryogenesis while the zygotic genome remains transcriptionally quiescent until the maternal-to-zygotic transition (MZT). Two interdependent events constitute the MZT in animals: the degradation of inherited maternal gene products and zygotic genome activation (ZGA) when the zygotic genome breaks transcriptional quiescence to produce transcripts that instruct subsequent embryogenesis. While the MZT is universal in multiple species, the underlying mechanisms, scale and timing of ZGA are diverse (reviewed in (Baroux *et al.*, 2008; Lee *et al.*, 2014; Tadros and Lipshitz, 2009; Walser and Lipshitz, 2011). Investigating how different species evolved various mechanisms to initiate ZGA is crucial to understanding embryogenesis.

Compared to animals, the timing and requirements of ZGA in flowering plants is limited. Histological and molecular evidence in *Hyacinthus orientalis* (hyacinth, (Niedojadło *et al.*, 2012; Pięciński *et al.*, 2008), *Nicotiana tabacum* (tobacco, (Ning *et al.*, 2006; Zhao *et al.*, 2011), *Oryza sativa* (rice, (Abiko *et al.*, 2013; Anderson *et al.*, 2013; Anderson *et al.*, 2017; Ohnishi *et al.*, 2014; Ohnishi *et al.*, 2019), *Triticum aestivum* (wheat, (Domoki *et al.*, 2013; Sprunck *et al.*, 2005) and *Zea mays* (maize, (Chen *et al.*, 2017; Dresselhaus *et al.*, 1999; Meyer and Scholten, 2007; Okamoto *et al.*, 2005; Sauter *et al.*, 1998) altogether indicate that large-scale transcriptional activities increase in zygotes after fertilization and prior to the first division. These results suggest that, similar to animals, plant zygotic genomes may also transition from a transcriptionally quiescent to active state. However, plant and animal life cycles are fundamentally different, where plants alternate between haploid gametophytic and diploid sporophytic phases (Walbot and Evans, 2003). More specifically, a subset of sporophytic cells undergo meiosis to produce haploid spores, which divide mitotically to generate multicellular gametophytes. The fertilization of egg cells contained within female gametophytes marks the onset of the sporophytic generation. Although it is unclear how similar the gametophytic-to-sporophytic transition in plants is to the MZT in animals, we have referred to the large-scale increase of transcriptional activities upon fertilization as ZGA below.

Although ZGA has been partially characterized in the model flowering plant *Arabidopsis thaliana* (Arabidopsis), the timing, parental contributions and requirements of ZGA was debatable. One model proposed that Arabidopsis zygotes are transcriptionally quiescent (Pillot *et al.*, 2010) and early embryos mostly rely on maternal gene products for growth and division (Del Toro-De León *et al.*, 2014; Vielle-Calzada *et al.*, 2000; Autran *et al.*, 2011; García-Aguilar and Gillmor, 2015; Armenta-Medina and Gillmor, 2019). However, several mutants exhibiting defects in the initial asymmetric division of the zygote segregate in a recessive manner consistent with transcriptional activities of either parental allele being sufficient for the first zygotic division (Yu *et al.*, 2016; Guo *et al.*, 2016; Xu *et al.*, 2005; Arnaud Ronceret *et al.*, 2008; A. Ronceret *et al.*, 2008; Ronceret *et al.*, 2005; Lin *et al.*, 2007; Liu and Meinke, 1998). Moreover, transcriptome analyses indicated equal parental genomic contributions to the embryonic transcriptome as early as the 1-cell/2-cell stage (Nodine and Bartel, 2012). Based on these results, it was proposed that the zygotic genome is activated within the first few hours after fertilization with equal contributions of maternal and paternal alleles to the transcriptome (Nodine and Bartel, 2012). Although the maternal transcriptome dominance reported in a conflicting report (Autran *et al.*, 2011) can be readily explained by the amount of maternal RNA contamination in the samples (Schon and Nodine, 2017), the precise timing and requirements of zygotic genome activation was unresolved until recently (Zhao *et al.*, 2019). Here, we provide independent evidence that transcriptional activities are markedly increased upon fertilization in Arabidopsis and that zygotic transcription is essential for the initial embryonic cell divisions.

## RESULTS

### Expansion Microscopy Improves Whole-Mount Fluorescence Immunostaining

Phosphorylated serine 2 of the carboxy-terminal domain of RNAPII (RNAPII Ser2P) indicates elongating polymerase (Hajheidari *et al.*, 2013). Therefore, we used conventional whole-mount fluorescence immunostaining (Pillot *et al.*, 2010; García-Aguilar and Autran, 2018) on fertilized ovules (seeds) to detect RNAPII transcriptional activities in zygotes and embryos. We also stained against tubulin with antibodies and chromatin with 4’,6-diamidino-2-phenylindole (DAPI) to unambiguously identify egg and zygote nuclei. We obtained several samples with consistent and strong signals, but found that the conventional protocol produced inconsistent results (Figure 1A). Namely, 92/234 (39%) samples exhibited uneven or no signal likely due to the limited antibody accessibility (Figure 1B). Embryos in particular had weak signals because they are embedded within seeds. Moreover, 77/234 (33%) samples had collapsed embryo sacs and were impossible to analyze (Figure 1B). We therefore could not robustly detect RNAPII Ser2P with the conventional immunostaining protocol.

**Figure 1.**
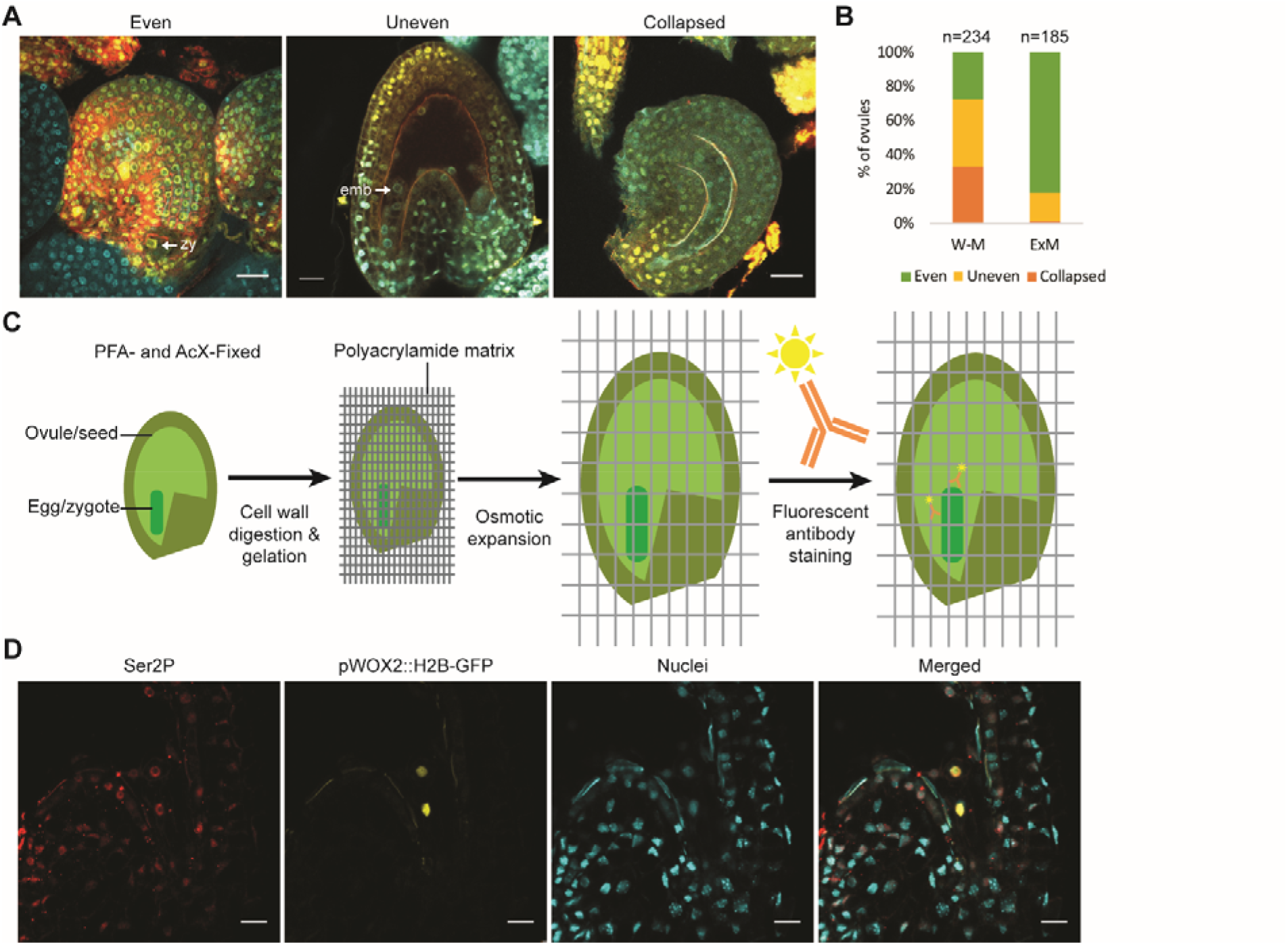
Expansion Microscopy on Arabidopsis Seeds. **(A)** Representative images of evenly and unevenly stained samples, or collapsed samples. Tubulin (red) and RNAPII Ser2P (yellow) were detected with immunofluorescence, and nuclei were stained with DAPI (cyan). **(B)** Quantification of the number of seeds with either even or uneven staining, or that were collapsed when using conventional whole-mount (W-H) or expansion microscopy (ExM) protocols. The total number of seeds examined with each method is indicated. **(C)** Schematic of expansion microscopy (ExM) workflow. Ovules/seeds were fixed, incubated with cell wall digestion enzymes, embedded in a polyacrylamide gel matrix and osmotically expanded before fluorescent immunostaining (See Materials and Methods for details). **(D)** Protein retention after expansion in embryos. Immunofluorescent signal from RNAPII Ser2P (red) or HISTONE 2B-tagged GFP expressed with the embryo-specific WOX2 promoter (pWOX2::H2B-GFP; yellow) are shown, as well as DAPI-stained nuclei (cyan). Scale bars represent 20 μm. zy, zygote; emb, embryo.

To improve the whole-mount fluorescent immunostaining method for Arabidopsis ovules/seeds, we adapted an expansion microscopy protocol (ExM, (Chen *et al.*, 2015; Tillberg *et al.*, 2016). The ExM technique physically expands specimen uniformly in three dimensions to increase reagent accessibility and microscopic resolution while retaining the relative position of signals. Samples were fixed in 4% paraformaldehyde and 0.1 mg/mL Acryloyl-X, SE (6- ((acryloyl)amino)hexanoic acid, succinimidyl ester) (AcX), incubated with cell wall-digesting enzymes and embedded in an expandable polyacrylamide gel matrix (Figure 1C). Samples were then expanded with osmotic pressure to improve antibody penetration and increase specimen size before immunostaining. We examined protein retention in expanded samples in seeds expressing a previously described embryo-specific reporter line (pWOX2::H2B-GFP, pWOX2::tdTomato-RCI2b; (Gooh *et al.*, 2015)). As expected, the H2B-GFP signal was confined to embryonic nuclei indicating that our protocol successfully retained proteins in their proper subcellular localizations (Figure 1D). The membrane-localized tdTomato-RCI2b signal was not observed demonstrating that the permeabilization removed membrane compartments and improved antibody penetration (Figure 1D). We also stained against RNAPII Ser2P and found that the epitopes were retained and detectable with antibodies (Figure 1D). To examine expansion ratios in three dimensions, we stained cell walls with SCRI Renaissance 2200 and nuclei with DAPI, and examined samples before and after expansion. Although expansions were not uniform in all three dimensions, ranging from 1.3 to 2.0 times in length depending on the experiment, the RNAPII Ser2P signal was strong and consistent among samples. With the ExM technique we found that only 2/185 (0.1%) samples had collapsed embryo sacs and 31/185 (16.7%) had uneven staining, while 152/185 (82.2%) samples had strong and consistent signal (Figure 1B). By physically expanding the specimens with the modified ExM technique before staining, we were therefore able to make interior tissues more accessible to reagents. This method produced more consistent staining and enabled the robust detection of subcellular marks within zygotes.

### Visualization of Transcriptional Activities in Eggs and Zygotes

We then employed the ExM approach described above to visualize RNAPII Ser2P in eggs and zygotes. We also stained against tubulin with antibodies and chromatin with DAPI to identify egg and zygote nuclei. In unfertilized ovules, the RNAPII Ser2P signal was weak in most (22/26, 85%) egg nuclei but strong in surrounding sporophytic ovule nuclei (Figure 2A). After fertilization, the RNAPII Ser2P signal was strong in most (27/29, 93%) zygote nuclei, as well as in surrounding endosperm and seed nuclei (Figure 2A).

**Figure 2.**
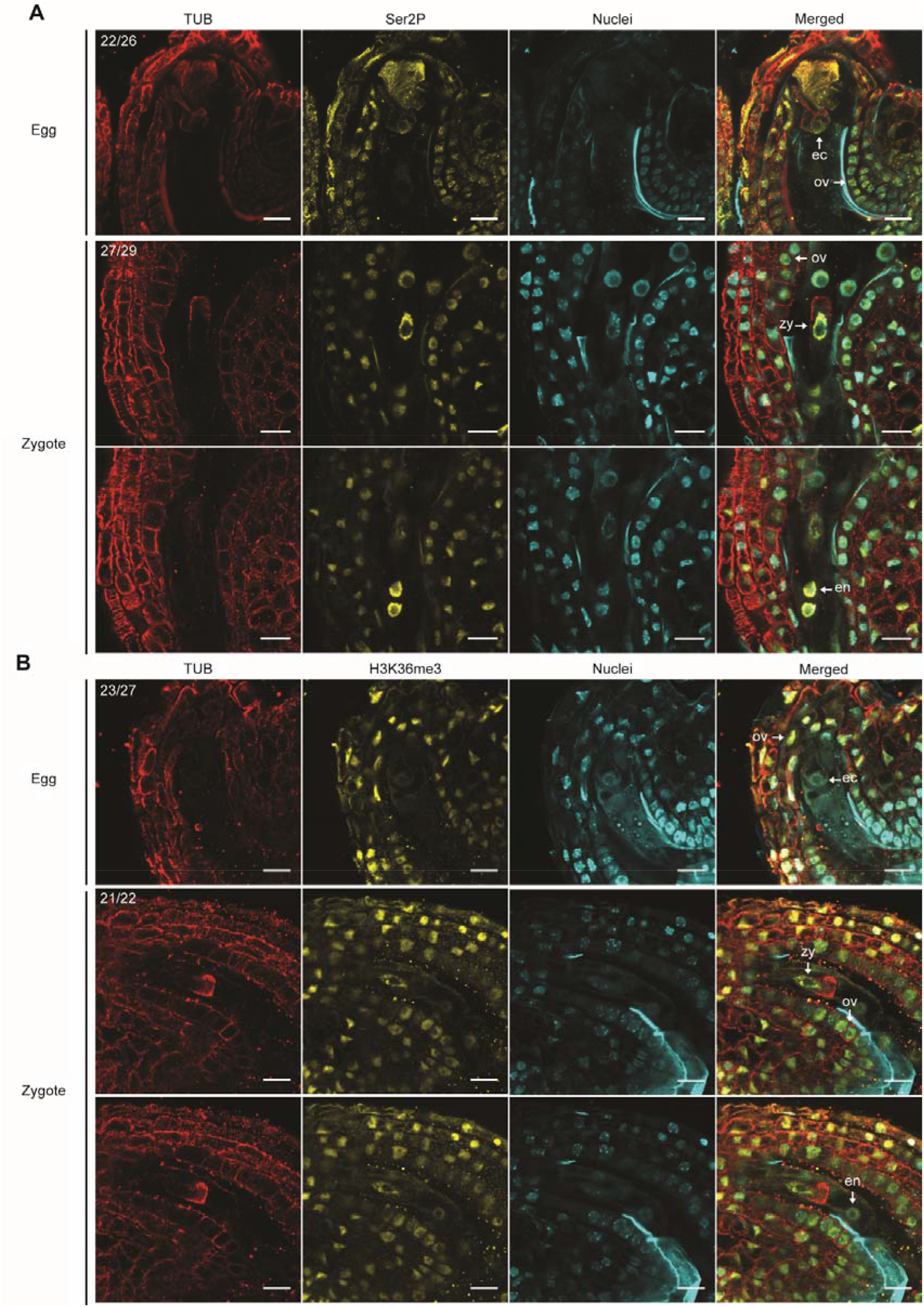
Visualization of Transcription in Eggs and Zygotes. **(A)** Representative ExM images of tubulin (red), RNAPII Ser2P (yellow) and DAPI-stained nuclei (cyan) in ovules containing eggs (*top*) or seeds containing zygotes with focal plane on zygotes (*middle*) or (Figure 2 continue) endosperm (*bottom*). The number of ovules or seeds with similar staining patterns out of the total number examined are indicated. **(B)** Representative ExM images of tubulin (red), H3K36me3 (yellow) and nuclei (cyan) in ovules/seeds containing eggs (*top*) or zygotes with focal planes on zygotes (*middle*) or endosperm (*bottom*). The number of ovules or seeds with similar staining patterns out of the total number of examined are indicated. Scale bars represent 20 μm. ec, egg cell; zy, zygote; en, endosperm; ov, ovule/seed tissue.

Histone H3 lysine 36 trimethylation (H3K36me3) is deposited during RNAPII transcription elongation (Wagner and Carpenter, 2012). Therefore, we examined H3K36me3 levels in ovules and seeds using ExM to further inspect transcriptional activities in eggs and zygotes, respectively. Similar to RNAPII Ser2P, H3K36me3 signal was weak in most (23/27, 85%) egg nuclei but strong in the surrounding sporophytic ovule nuclei (Figure 2B). In contrast, H3K36me3 signal was strong in most (21/22, 95%) zygote nuclei, as well as in endosperm and surrounding sporophytic ovule nuclei (Figure 2B). The detection of H3K36me3 and RNAPII Ser2P in eggs and zygotes indicated that RNAPII transcription was relatively low in eggs and became highly active in zygotes after fertilization.

### Active Transcription Is Physiologically Required for Zygote Cell Division

Classic transcriptional inhibition experiments in mouse (Braude *et al.*, 1979), *Drosophila melanogaster* (Edgar and Datar, 1996), *Caenorhabditis elegans (Edgar et al., 1994), Xenopus laevis* (Newport and Kirschner, 1982) and *Danio rerio* (Zamir *et al.*, 1997) early embryos elegantly demonstrated that inherited gene products were sufficient for early embryogenesis. Therefore, to test whether active transcription is required for Arabidopsis zygote development, we cultured seeds in the presence of RNAPII inhibitors and examined zygotic cell division patterns with live-imaging microscopy (Gooh *et al.*, 2015). If inherited gene products were sufficient for early embryogenesis, then we expected no developmental delay or arrest when transcriptional inhibitors were included in the culture media. Seeds were cultured in the presence or absence of RNAPII inhibitors for one hour before acquiring the first image, and zygotes were labelled with nuclear-localized GFP and plasma membrane-localized tdTomato both of which were under the control of the embryo-specific WOX2 promoter (pWOX2::H2B-GFP, pWOX2::tdTomato-RCI2b; (Gooh *et al.*, 2015). Similar to what was originally reported for this ovule culture system, we found that approximately 80% of embryos survived in our system under all conditions (Figure 3G). That is, regardless of culture conditions about 20% of embryos lost fluorescent signal likely due to technical limitations of the system. We classified these embryos as dead and the remaining ones as survived, and only the surviving embryos were informative to test our hypothesis. As a positive control, we first compared cell division patterns of embryos cultured with or without the flavopiridol (FLP) kinase inhibitor, which causes cell-cycle arrest (Dai and Grant, 2003). As expected, all 28 surviving embryos cultured with 100 μM FLP had arrested cell division whereas in N5T control media all surviving embryos divided normally (Figures 3A and 3B, Movie S1 and S2).

**Figure 3.**
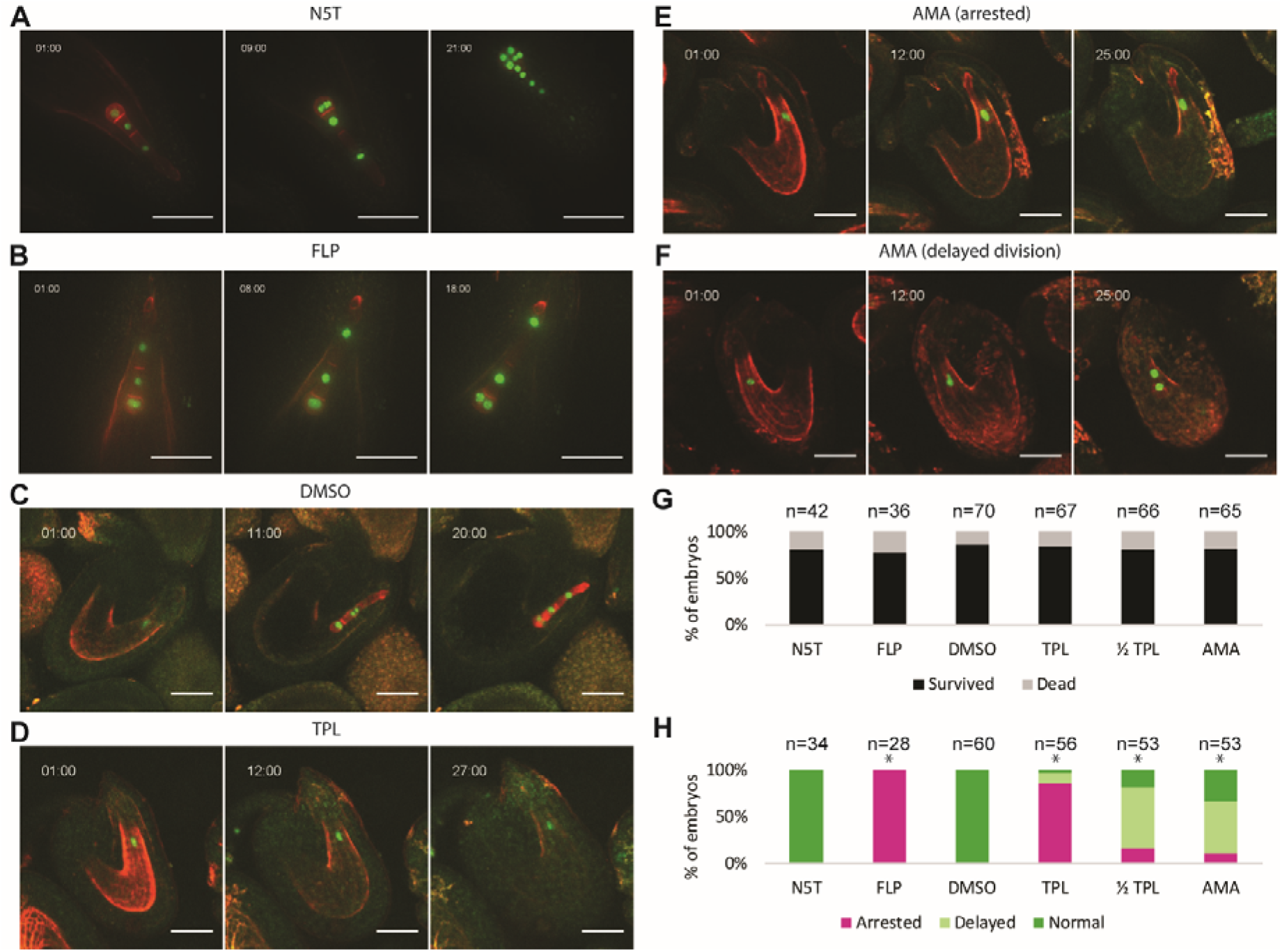
Transcription Is Required for Initial Embryonic Cell Divisions. **(A - F)** Time-lapse observations of embryos expressing pWOX2::H2B-GFP, pWOX2::tdTomato-RCI2b cultured in N5T **(A)**, N5T with 100 μM FLP **(B)**, N5T with 0.5% DMSO **(C)**, N5T with 0.5% DMSO and 500 μM TPL **(D)**, or N5T with 0.5% DMSO and 100 μg/mL AMA **(E,F)**. The time after incubation is indicated as hours:minutes. **(G)** Bar chart illustrating the percentage of embryos that survived or died under various conditions. The total number of embryos examined for each condition are shown. **(H)** Bar chart showing the percentage of embryos that divided normally or abnormally when cultured under various conditions. The total number of surviving embryos for each condition are indicated. Asterisks indicate significantly different population determined by chi-square tests, *P*-values < 3.6 × 10^-8^. Scale bars represent 50 μm.

Similar to N5T control media, all surviving embryos within seeds cultured with 0.5% dimethyl sulfoxide (DMSO) divided normally with approximately six hours intervals between each cell division (n = 60, Figure 3C and Movie S3; *P*-value = 0.69, chi-square test). Triptolide (TPL) induces protease-dependent degradation of RNAPII components (Titov *et al.*, 2011; Vispé *et al.*, 2009), and we cultured seeds with TPL to test whether active transcription was required for zygotic divisions. Fifty-seven embryos survived when cultured with 500 μM TPL in 0.5% DMSO, and of these 48 (84.2%) were arrested at the zygote stage, 6 (10.5%) had delayed cell divisions (>12 hours between cell divisions) and only 2 (3.5%) developed normally (Figure 3D and Movie S4). The proportion of embryos that were arrested or delayed was significantly greater than observed for DMSO control condition (*P*-value = 2.2 × 10^-16^, chi-square test). Moreover, TPL had a dose-dependent effect on embryo development whereupon when seeds were cultured with 250 μM TPL, 9 out of the 53 (16.9%) surviving zygotes were arrested, 34 (64.2%) had delayed divisions and 10 (18.9%) developed normally (*P*-value = 3.3 × 10^-13^ compared to DMSO control, chi-square test).

To further test whether de novo transcription was required for zygote divisions, we also cultured seeds with α-amanitin (AMA), which binds to RNAPII and prevents nucleotide incorporation and transcript translocation (Kaplan *et al.*, 2008; Brueckner and Cramer, 2008). In the presence of 100 μg/mL AMA, 53 zygotes survived the culture and of these only 18 (34.0%) divided normally, while 6 (11.3%) were arrested (Figure 3E and Movie S5) and 29 (54.7%) were delayed (Figure 3F and Movie S6) in their development (*P*-value = 3.6 × 10^-8^ compared to DMSO control, chi-square test). The arrested and delayed zygote divisions in the presence of TPL and AMA transcription inhibitors indicated that de novo transcription was essential for the onset of zygote division.

## DISCUSSION

Optimization of the ExM fluorescent immunostaining technique for seeds enabled the robust detection of RNAPII Ser2P and H3K36me3 in zygotes. Because RNAPII Ser2P and H3K36me3 are hallmarks of active transcription, this indicated that zygotes are transcriptionally active soon after fertilization and before the first division. Moreover, zygotes had arrested and delayed cell divisions when cultured in the presence of transcriptional inhibitors (i.e. TPL and AMA), which is consistent with ZGA being required for the first division. Our observations complement a recent transcriptome study demonstrating that ZGA occurs in zygotes and is required for elongation and division (Zhao *et al.*, 2019).

While the conventional whole-mount immunostaining protocol can result in even-stained samples, we found that the method produced variable and inconsistent results. To improve the robustness of histological detection in Arabidopsis ovules/seeds, we adapted ExM to plant tissues in this study. Constraints imposed by undigested cytoskeleton and cell wall components limited the ability to fully expand samples. We omitted the proteinase K treatment described in the original ExM protocol because we found that even a mild proteinase K treatment (i.e. 5 U/mL for 30 minutes) resulted in signal loss without improving expansion ratios or consistency. Nevertheless, we were able to produce expanded samples with the ExM method, which made interior tissues accessible to antibodies and resulted in more consistent immunostaining. Sample expansion also helped reduce autofluorescence, which is common in plant tissues. Our ExM protocol therefore provides a more robust and efficient option for conducting whole-mount fluorescent immunostaining of ovules/seeds including those containing zygotes.

The detection of weak levels of RNAPII Ser2P and H3K36me3 in mature eggs indicated that there is low transcriptional activity before fertilization. After fertilization, signal corresponding to RNAPII Ser2P and H3K36me3 was strong demonstrating that transcriptional activities dramatically increased in the zygote before the first division. It was previously reported that endosperm nuclei but not in zygote nuclei were transcriptionally active (Pillot *et al.*, 2010). However, our observations indicate that both endosperm and zygote nuclei are transcriptionally active. The previous inability to detect RNAPII Ser2P signal in zygotes may be due to the antibodies used (H5, ab24758; Abcam, discontinued) or the variability of the conventional whole-mount protocol which is especially problematic for the deeply embedded zygotes.

As revealed by live-cell imaging, transcriptional inhibition resulted in delayed or arrested development and provided evidence that ZGA is required for initial zygotic growth and division. That is, treatment with the TPL inhibitor consistently arrested zygotes while AMA treatment at least doubled the time required for cell division. The different responses of zygotes to TPL and AMA were likely due to the incubation timing because TPL and AMA are known to be fast-response and slow-response inhibitors, respectively (Bensaude, 2011). The 250 μM TPL treatment, which is at least 100-fold greater than the recommended concentration for tissue culture (Bensaude, 2011), showed delayed instead of arrested zygote division. The requirement of high concentrations of transcriptional inhibitors to have a physiological effect may be due to the high transcriptional activities of zygotes. Consistent with our results using transcriptional inhibitors, previous RNAi-mediated knock-down of RNAPII resulted in delayed embryogenesis (Pillot *et al.*, 2010). Although we cannot completely exclude that the transcriptional inhibition of ovule tissues supporting the developing embryo (e.g. integuments and endosperm) are the primary cause of the embryo arrest we observed, it is clear that inherited parental transcripts are not sufficient and de novo transcripts are required for early embryogenesis regardless of their origin. Moreover, in vitro fertilized or isolated zygotes developed normally through early embryogenesis in the absence of surrounding maternal tissue in rice (Sato *et al.*, 2010; Uchiumi *et al.*, 2007), maize (Kranz and Lorz, 1993) and tobacco (Zhao *et al.*, 2011; He *et al.*, 2007), and Arabidopsis embryos can develop at least to the globular stage without endosperm (Gooh *et al.*, 2015; Ngo *et al.*, 2007). These reports are consistent with the delayed and arrested zygotic division observed upon culturing with transcriptional inhibitors being primarily caused by the inability to transcribe genes in zygotes.

Because it is difficult to differentiate de novo transcribed from maternally inherited transcripts, the activation of the zygotic genome can be inferred from the transcriptional activities of paternal alleles. Several studies reported that early Arabidopsis embryos rely on maternal factors with little or no paternal activity (Golden *et al.*, 2002; Del Toro-De León *et al.*, 2014; Vielle-Calzada *et al.*, 2000; Grossniklaus *et al.*, 1998; Luo *et al.*, 2000; Guitton and Berger, 2005; Pagnussat *et al.*, 2005) and suggest a quiescent state in zygotes (Pillot *et al.*, 2010). In contrast, other studies reported paternal allele expression in early embryos (Aw *et al.*, 2010; Xiang *et al.*, 2011; Köhler *et al.*, 2005; Weijers *et al.*, 2001; Baroux *et al.*, 2001; Ueda *et al.*, 2011). Additionally, mutants defective in the asymmetric division of the zygote were reported to be recessive, suggesting that both parental alleles are transcriptionally active after fertilization (Lukowitz *et al.*, 2004; Yu *et al.*, 2016; Guo *et al.*, 2016; Xu *et al.*, 2005; Ronceret *et al.*, 2005; A. Ronceret *et al.*, 2008; Arnaud Ronceret *et al.*, 2008; Lin *et al.*, 2007; Liu and Meinke, 1998; Tzafrir *et al.*, 2004). Moreover, a recent genome-wide study reported significant upregulation of 4,436 genes in zygotes compared to eggs (Zhao *et al.*, 2019). Altogether these results indicate that genes involved in early embryogenesis are transcriptionally active in zygotes, and thus do not support gradual ZGA or reliance on maternal gene products during early Arabidopsis embryogenesis as previously proposed (García-Aguilar and Gillmor, 2015; Armenta-Medina and Gillmor, 2019).

In addition to Arabidopsis, genetic, microscopic and genomic studies in multiple flowering plants, including maize (Chen *et al.*, 2017; Dresselhaus *et al.*, 1999; Meyer and Scholten, 2007; Okamoto *et al.*, 2005; Sauter *et al.*, 1998; Scholten *et al.*, 2002), wheat ((Domoki *et al.*, 2013; Sprunck *et al.*, 2005)), tobacco (Ning *et al.*, 2006; Zhao *et al.*, 2011; Xin *et al.*, 2011) and rice ((Abiko *et al.*, 2013; Anderson *et al.*, 2013; Anderson *et al.*, 2017; Ohnishi *et al.*, 2014; Ohnishi *et al.*, 2019)), indicate that ZGA occurs soon after fertilization in zygotes. In plants, zygotes mark the transition from the haploid gametophytic to diploid sporophytic phase of the life cycle. Because there is no clear evidence of prolonged transcriptional quiescence after fertilization and parentally inherited gene products are not sufficient for early embryogenesis, the transcriptome remodeling observed in plant zygotes during the gametophytic-to-sporophytic transition is fundamentally different than the maternal-to-zygotic transition in early animal embryos as previously proposed (Ueda *et al.*, 2017). In further contrast to animals (Tadros and Lipshitz, 2009; Lee *et al.*, 2014), the apparent similarities in the timing of transcriptome remodeling across plant species is intriguing and may indicate similar underlying mechanisms shared among different plant species. For example, maternally and paternally inherited gene products converge to rapidly activate *WUS HOMEOBOX8 (WOX8)* gene expression in Arabidopsis zygotes (Lukowitz *et al.*, 2004; Bayer *et al.*, 2009; Ueda *et al.*, 2011; Ueda *et al.*, 2017), and similar mechanisms integrating biparentally inherited information may exist throughout Arabidopsis and other plant genomes. With published egg and zygote transcriptomes in multiple plants, as well as the increasing number of genome-wide approaches applicable to low amounts of input material, the field is poised to identify the key regulatory genes involved in transcriptome remodeling and initiation of embryogenesis.

## MATERIALS AND METHODS

### Plant Materials and Growth Conditions

*Arabidopsis thaliana* accession Columbia (Col-0) and pWOX2::H2B-GFP, pWOX2::tdTomato-RCI2b transgenic Col-0 plants were grown in a climate-controlled growth chamber with 20°C-22°C temperature and 16h light/8h dark cycles.

### Cell Wall Digestion Enzymes

We tested several cell wall digestion enzymes from multiple manufacturers and found the performance varied between manufacturers as well as between batches. We tested driselase (Sigma), cellulase (Sigma), cellulase R10 (Duchefa, Yakult), cellulase RS (Duchefa, Yakult), pectolyase (Duchefa, discontinued), pectinase (Sigma), macerozyme R10 (Duchefa) and hemicellulase (Sigma) for conventional whole-mount protocol. We chose cellulase RS (Duchefa), hemicellulase (Sigma) and pectinase (Sigma) or macerozyme R10 (Duchefa) for expansion microscopy based on their performance and availability.

### Conventional Whole-Mount Fluorescent Immunostaining

Conventional whole-mount fluorescent immunostaining was performed according to a published protocol (García-Aguilar and Autran, 2018). Seeds of self-fertilized Col-0 siliques at stages 14-15 (Smyth *et al.*, 1990) were isolated under a dissection scope. Isolated seeds were collected in 4% PFA, 0.1% Triton X-100 and 1 × PBS solution. Seeds were then briefly vacuum infiltrated and incubated at room temperature for one hour. Fixed seeds were washed three times with 0.1% Triton X-100 and 1 × PBS before incubation with enzyme mix (1% driselase, 0.5% cellulase, and 1% pectolyase in water). Alternatively, seeds were washed once more with protoplast salt solution (20 mM MES, pH 5.0, 0.4 M mannitol, 20 mM KCl and 10 mM CaCl_2_) before incubation with protoplast enzyme solution (3% cellulase, 1% hemicellulase, 1% macerozyme, 20 mM MES, pH5.0, 0.4 M mannitol, 20 mM KCl and 10 mM CaCl2, 0.1% BSA and 1% β-mercaptoethanol) before use. Enzyme solution was prepared as previously described (Yoo *et al*., 2007). Seeds were incubated in either enzyme solution at 37°C for 2 hours with gentle agitation. Digested seeds were washed twice with 0.2% Triton X-100, 1× PBS and embedded in 3% polyacrylamide matrix on adhesive slides as described (García-Aguilar and Autran, 2018). Slides were incubated with 1% Triton X-100, 1 × PBS at 4°C for two hours with gentle agitation. Permeabilized samples were incubated with 1 × PBS, 2% BSA, 0.1% Triton X-100 at room temperature for one hour for blocking. Samples were then incubated with 1 × PBS, 2% BSA, 0.1% Triton X-100 and 1:500 dilution of primary antibodies against RNAPII Ser2P (ab5095, Abcam) and tubulin (ab89984, Abcam) at 4°C overnight with gentle agitation. On the next day samples were washed with 0.2% Triton X-100, 1× PBS at 4°C for one hour at least five times. For secondary antibody incubation, samples were incubated in 1:500 dilution of anti-rabbit-Alexa488 (ab150077, Abcam) and anti-chicken-Alexa555 (ab150170, Abcam) in 1× PBS, 2% BSA, 0.1% Triton X-100 at 4°C overnight with gentle agitation. Samples were then washed with 10 μg/mL 4’,6-diamidino-2-phenylindole (DAPI), 0.2% Triton X-100, 1× PBS at 4°C in the dark for one hour twice and washed three times without DAPI. Samples were mounted in Vectashield Mounting Medium (H-1200, Vector) and imaged by ZEISS LSM700/780 with 25× oil objective. Color channels were scanned sequentially to avoid false signal.

### Expansion Microscopy

We modified published ExM protocols for plant tissue (Chen *et al.*, 2015; Tillberg *et al.*, 2016). Pistils or siliques were carefully sliced open longitudinally under a dissection scope and transferred to 1 × PBS, 0.1% Triton X-100, 4% PFA and 0.1 mg/mL Acryloyl-X, SE (6- ((acryloyl)amino)hexanoic acid, succinimidyl ester; Thermo Fisher). After brief vacuum infiltration, samples were incubated at 4°C overnight. Samples were then washed with water twice and washed once more with protoplast salt solution (20 mM MES, pH5.0, 0.4 M mannitol, 20 mM KCl and 10 mM CaCl2) before incubated with protoplast enzyme solution (3% cellulase, 1% hemicellulase, 1% macerozyme, 20 mM MES, pH 5.0, 0.4 M mannitol, 20 mM KCl and 10 mM CaCl2, 0.1% BSA and 1% β-mercaptoethanol before use. (Yoo *et al.*, 2007). Samples were incubated at 37°C for 2-3 hours (depending on developmental stage) with gentle agitation. Cell wall-digested samples were washed twice with 0.2% Triton X-100, 1 × PBS and then permeabilized with 1% Triton X-100, 1× PBS at 4°C for two hours. Pistils or siliques were then carefully dissected on depression slides under a dissection scope. Strings of ovules/seeds attached to septums (i.e. ovule strings) were detached from pistils or siliques and transferred to 0.2% Triton X-100, 1 × PBS. Isolated ovule strings were drained briefly and incubated in monomer solution (1× PBS, 2 M NaCl, 8.625% (w/w) sodium acrylate, 2.5% (w/w) acrylamide, 0.15% (w/w) N,N’-methylenebisacrylamide) at 4°C overnight. Samples were then polymerized with 0.2% ammonium persulfate (APS) and tetramethylethylenediamine (TEMED) on glass slides with an adequate spacer (200-300 μm) at room temperature for one hour. Proteinase K treatment was omitted because it resulted in massive loss of epitopes. Gel slices containing ovule strings were cut out with razor blades and every two slices were transferred to 1 mL water in 2 mL microtubes for expansion for one hour. The expansion was repeated at least twice. Expanded samples were then immunostained and imaged as described above.

### Live-Cell Imaging

The ovule culture was performed as previously reported with slight modifications (Gooh *et al.*, 2015). Seeds from self-pollinated flowers at stage 14-15 (Smyth *et al.*, 1990) were carefully collected under a dissection scope and transferred to N5T medium (5% trehalose dihydrate, 1 × Nitsch basal salt mixture, 1 × Gamborg’s vitamin solution, 0.05% MES-KOH, pH 5.8). A brief vacuum infiltration was applied to seeds followed by a 30-minute incubation in N5T medium or medium supplemented with α-amanitin, flavopiridol, triptolide, or dimethyl sulfoxide at room temperature with gentle agitation to submerge seeds completely. The seeds were then transferred to micro-Insert 4 Well in μ-Dish (ibidi) with corresponding medium for live-cell imaging with either Yokogawa CSU X1 spinning disc and Axio Observer (ZEISS) or Visiscope Spinning Disc Confocal (Visitron Systems GmbH). The first images were taken about one hour after incubation and the time-lapsed images were taken every 30 to 60 minutes for at least 20 hours.

### Image Processing

All confocal microscope images were adjusted by ZEN software before exporting and scale bars were added by Fiji. Spinning disc microscope images were processed by Fiji. For each timepoint the Z-stacks were merged by maximum intensity projection, and then the contrast was adjusted for each channel. Color channels were then merged and images were cropped before adding scale bars and time stamps.

## ACKNOWLEDGEMENTS

This work was supported by the Österreichische Akademie der Wissenschaften (Austrian Academy of Sciences). Special thanks to Daisuke Kurihara and Tetsuya Higashiyama for generously sharing pWOX2::H2B-GFP, pWOX2::tdTomato-RCI2b lines and their instructions on live-cell imaging. We are also grateful to Frederic Berger for insightful discussions and Michael Schon for his input on figure generation. Additionally, we acknowledge the IMP-IMBA-GMI BioOptics Core Facility and the Plant Sciences Facility at Vienna BioCenter Core Facilities GmbH (VBCF) for support. Ping Kao also personally thanks Kotoha Tanaka and Kaori Sakuramori for their support.

## DECLARATION OF INTERESTS

The authors declare they have no conflict of interest.

## AUTHOR CONTRIBUTIONS

Conceptualization, P.K. and M.D.N.; Methodology, P.K.; Formal Analysis, P.K.; Investigation, P.K.; Writing – Original Draft, P.K.; Writing – Review & Editing, M.D.N. and P.K.; Visualization, P.K.; Supervision, M.N.

## LEGENDS FOR SUPPORTING INFORMATION

**Movie S1.** Live-cell imaging of embryos cultured in N5T medium.

**Movie S2.** Live-cell imaging of embryos cultured with 100 μM FLP in N5T medium.

**Movie S3.** Live-cell imaging of embryos cultured with 0.5% DMSO in N5T medium.

**Movie S4.** Live-cell imaging of embryos cultured with 500 μM TPL, 0.5% DMSO in N5T medium.

**Movie S5.** Live-cell imaging of embryos cultured with 100 μg/mL AMA, 0.5% DMSO in N5T medium with arrested zygotes.

**Movie S6.** Live-cell imaging of embryos cultured with 100 μg/mL AMA, 0.5% DMSO in N5T medium with delayed division.

